# Representation learning of single-cell time-series with deep variational autoencoders

**DOI:** 10.1101/2025.09.22.677729

**Authors:** Achille Fraisse, Diego A. Oyarzún, Meriem El Karoui

## Abstract

Single-cell technologies have led to many insights on the function of individual cells within populations. Time-series data, in particular, have become increasingly adopted to uncover the molecular basis of heterogeneity in processes such as gene regulation, growth adaptation, and drug resistance. Yet the analysis of single-cell trajectories often requires manual and application-dependent techniques to extract meaningful features that correlate with the biological processes of interest. Here, we employ representation learning to automatically encode time-series into a low-dimensional feature space that can be used for downstream prediction tasks. We trained deep variational autoencoders on cell length data from *Escherichia coli* cells growing in a “mother machine” device that allows tracking single cells over long periods. We show that the learned representations preserve the structure of the data across various growth media. Using the pretrained model in tandem with supervised learning, we encoded fresh time-series from cells exposed to single and combination antibiotics, which achieved excellent classification accuracy across drug treatments. Moreover, we demonstrate that the learned embeddings can be used to accurately classify datasets from different laboratories and growth conditions without retraining. This demonstrates that the autoencoder extracts meaningful information with promising generalization power. Our results highlight the potential of representation learning for the analysis of single-cell responses under chemical perturbations and growth conditions.

## I. INTRODUCTION

In the last three decades, single-cell analysis has emerged as an essential approach for understanding the phenotypic and functional diversity of individual cells in isogenic populations, which is often masked by bulk measurements. In bacteria, this approach has provided insights into the roles that heterogeneity plays in cellular processes such as gene expressio^1^, antibiotic resistance^2^, and stress responses^3^. Importantly, single-cell analysis is often conducted by tracking individual cells over time, thereby revealing not only heterogeneity within a population but also the dynamic nature of this variation. The quantification of such cellto-cell variation often depends on analysing a large number of time-series and determining correlations with observed phenotypes or experimental conditions. This requires algorithms that can reliably process the data and extract features for downstream analyses such as discovering subpopulations or predicting phenotypic endpoints.

Traditionally, feature extraction in biological time-series involves manual selection and computation of statistical features from the data. Numerous tools have been developed to this end that allow for computation of a wide range of time-series descriptors^4,5^. These approaches, however, can be limited by their reliance on prior knowledge and intuition about which features are most relevant for a particular phenotype or downstream task. Moreover, manual feature extraction is application-dependent and may not capture all the complex and nonlinear relationships present in time-series data, especially in biological contexts where the underlying processes are only partially understood.

Advances in machine learning have complemented such approaches with methods for encoding time-series automatically. Specifically, representation learning can automate the feature extraction process by extracting meaningful information from the data itself^6^. This approach has been widely successful in computer vision, natural language processing and other domains, and has the potential to uncover hidden dependencies in the time-series that are not readily apparent through manual methods^7^. One strategy is the use of autoencoder algorithms that project time-series data into a low-dimensional latent space that can then be employed for clustering, classification and regression tasks^8,9^.

Here, we describe the use of representation learning for the analysis of single-cell timeseries data. We focus on data obtained from the “mother machine”^10–12^, a microfluidic device that traps individual bacterial cells in micro-channels and allows imaging of steady-state growth during long periods. These devices have enabled the collection of high-quality data from single cells in a growing number of biological questions, including bacterial ageing^10^, stochastic gene expressio^13^, cell size homeostasis^14,15^, antibiotic responses^16^, detection of mutations^17,18^, or detection of infectio^19^. Multiple tools exist to analyse the images resulting from these experiments and perform segmentation and tracking^20–22^. They produce a large number of time traces for cell length (a proxy for cell volume), the expression of fluorescent-tagged proteins or transcriptional reporters, and other phenotypes of interest.

We sought to assess the potential of representation learning to compress mother machine time-series data into a meaningful representation that can be used for various prediction tasks. To this end, we trained a Variational Recurrent Autoencoder (VRAE^23^) model on temporal data from *Escherichia coli* growing under various carbon sources in a mother machine. We focused on single-cell length data as this is a common readout affected by many factors, including growth conditions^24,25^ and chemical perturbations^26,27^, and is available in many mother machine experiments. Using clustering algorithms, the results demonstrate that the learned embeddings preserve the structure and separability of the data across growth conditions. We then encoded unseen time-series from cells treated with single and combination antibiotics^28^, and employed the embeddings for training accurate classifiers of drug treatment. To test the ability of the pretrained encoded to transfer to new datasets, we challenged the model with an external dataset obtained in a different lab and using different growth temperatures^29^. Without retraining, the model was able to encode the new timeseries into meaningful representations that allowed for clustering and classification of cells according to the temperature in which they were grown. Altogether, our results underscore the power and utility of representation learning for the analysis of mother machine data across multiple application domains.

## II. RESULTS

### A. Learning representations of single-cell time-series with a variational recurrent autoencoder

To learn representations of single cell trajectories from mother machine devices (Figure 1A), we employed a variational autoencoder (VAE) designed to compress cell length trajectories of single *Escherichia coli* cells growing in three media: glycerol (gly, slow growth rate), glucose (glu, medium growth rate), and glucose supplemented with amino acids (gluaa, fast growth rate). The VAE architecture is widely employed as a strategy to extract meaningful features from unlabelled data, and is based on learning the probability distribution of data when projected into a latent space^30^. Typical VAE architectures employ feedforward neural networks to encode the data into a latent representation, and then decode it into the original feature space. Both networks are jointly trained to minimize a loss function that accounts for the mismatch between the input and the reconstruction, as well as the distance between the distribution of samples in the original and the embedding space. This enables the VAE to learn a low-dimensional representation that preserves the structure of the original data. In the case of time-series data, however, to capture dependencies between time points, we opted for an architecture based on recurrent neural networks that is better suited for sequential data^8,31,32^. We adopted the Variational Recurrent Autoencoder (VRAE) model from Fabius and Amersfoort^23^, which employs Long Short-Term Memory (LSTM) networks^33^ as encoders and decoders, and allows capturing long-range dependencies between time points. Similar architectures based on LSTM units have been recently employed for processing mother machine data and optogenetic control of single cells^34^.

**FIG. 1.**
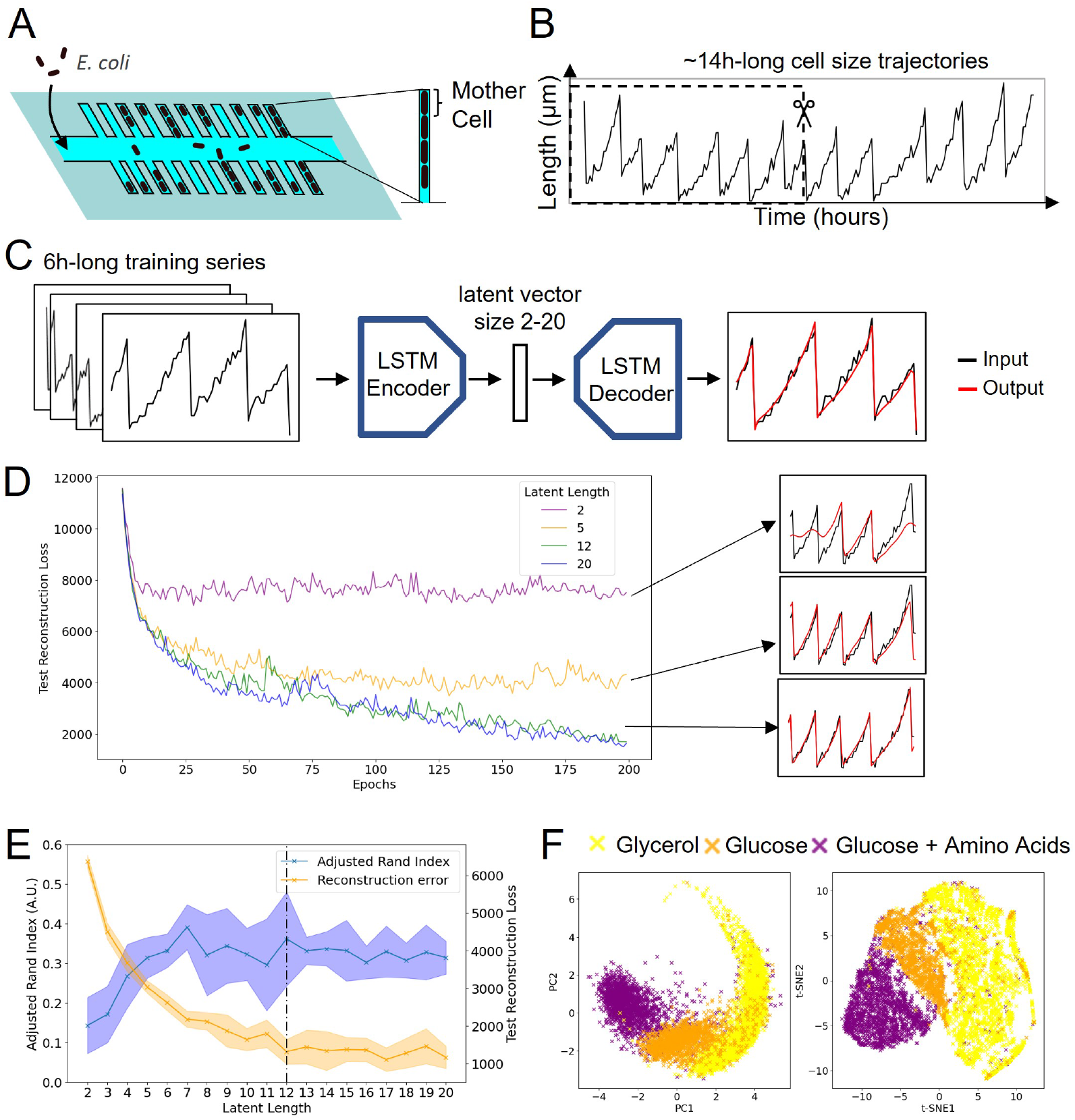
Training an Autoencoder on Single-Cell Time-series Data. **A**: *E. coli* cells grow and divide in the microchannels of a “Mother Machine” device10. **B**: By imaging every 5 or 10 minutes, depending on the media, the length of the first cell in the channel or “Mother cell” was tracked to form a time-series of cell size. **C**: time-series used for training are chopped into 6-hour fragments, using a rolling window of 30 minutes. A variational recurrent autoencoder is then trained to encode the time-series into a latent vector and then reconstruct it. **D**: Evolution of the test error (reconstruction error on the test set, mean over 10 trainings) during training for different latent vector sizes. On the right are examples of time-series reconstruction using trained models of different latent vector sizes. **E**: Evaluation of the 19 trained models on the full dataset containing cells growing in three different media. The reconstruction error and Adjusted Rand Index (ARI), here used as the model’s ability to separate the three growth conditions in the latent space (see section IV C for more information), are shown for different latent lengths, displaying the mean and standard deviation over 10 training runs. The vertical line shows the chosen latent length of 12. **F**: PCA and t-SNE representation of trajectories from control experiments after being embedded using the 12-latent-dimensions autoencoder, colored by growth media.

Given the periodic nature of the cell length trajectories (Figure 1B), we reasoned that a few cell divisions are sufficient to learn the structure of the data. We thus sliced the timeseries into 6-hour fragments using a rolling window. This allows capturing the periodicity of the signal and, at the same time, augmenting the number of time-series to train the autoencoder. In total, we trained the autoencoder on 57607 time-series corresponding to 1675 cells (see Table I). Initial training results (Figure 1C) show that the autoencoder can effectively reconstruct the cell trajectories with high accuracy and has a denoising effect, a well-known property of this model architecture^35^. A key parameter of the autoencoder model is the dimensionality of the latent space, as this dictates the number of trainable weights of the encoder and decoder LSTM units. To explore the impact of the dimension of the latent space on VRAE performance, we first trained the model using 2 to 20 latent variables and examined the evolution of the reconstruction loss computed on a held-out test set along the training epochs. The results (Figure 1D) show a clear tradeoff between data compression and reconstruction quality. In line with expectation, a higher dimensionality leads to better reconstruction performance due to the improved expressiveness of the model.

**TABLE I.** Number of series before and after processing used in each figure. Ctrl = Control (no antibiotic), Cip = ciprofloxacin, Tet = tetracycline, Cip+Tet = ciprofloxacin + tetracycline. *: sliced in overlapping time-series of 6h (rolling window of 30 min. **: sliced in non-overlapping consecutive time-series of 6h.)

|  | # of series<br>(variable length) | # of series after<br>processing (6h length) |
| --- | --- | --- |
| Fig1 train | 1675 | 57607* |
| Fig1 test | 187 | 699** |
| Fig 2 train Ctrl Vs Cip | 942 | 942 |
| Fig 2 test Ctrl Vs Cip | 236 | 236 |
| Fig 2 train Ctrl Vs Tet | 899 | 899 |
| Fig 2 test Ctrl Vs Tet | 225 | 225 |
| Fig 2 train Ctrl Vs Cip+Tet | 896 | 896 |
| Fig 2 test Ctrl Vs Cip+Tet | 225 | 225 |
| Fig 3 set | 279 | 2329** |

We optimized the dimensionality of the latent space by testing the ability of the encoder to preserve data structure across the growth conditions. We first computed embeddings for time-series from control experiments not used in training, and then checked the separation of the three growth conditions by clustering the latent representations with the *k−*means algorithm (*k* = 3 clusters). The results (Figure 1E) suggest that ARI plateaus above 7 dimensions, while the best reconstruction quality is achieved for 12 or more, as suggested by Figure 1D. We therefore selected 12 as the dimension of latent space for the remainder of this study, as it achieved a reasonable balance between reconstruction quality and data compression. Two-dimensional visualizations (PCA, t-SNE) of the 12-dimensional latent space show a clear separation of the three growth conditions along a continuous manifold (Figure 1F), indicating that the representation captures biologically relevant information.

### B. Identification of chemical perturbations using the pretrained autoencoder and labelled data

A key application of microfluidics devices, such as the mother machine, is to perform antibiotic susceptibility assays^36^, because sensitive bacteria change growth rate when they are exposed to antibiotics, whereas resistant isolates continue to grow normally. To test whether our autoencoder captures these effects, we created a dataset of labelled time-series from cells exposed to tetracycline (tet), ciprofloxacin (cip) or a combination of both (ciptet), as shown in Figure 2A. For each mother cell, we extracted a 6-hour time-series between 2 and 8 hours, which corresponds to the first 6 hours of antibiotic exposure. The time-series were then rescaled and embedded with the autoencoder that had been previously trained on control data (with 12 latent variables, as in Figure 1F). This allowed us to test the generalisation power of the learned embeddings on a fresh dataset with time-series and labels not seen during training (Figure 2B).

**FIG. 2.**
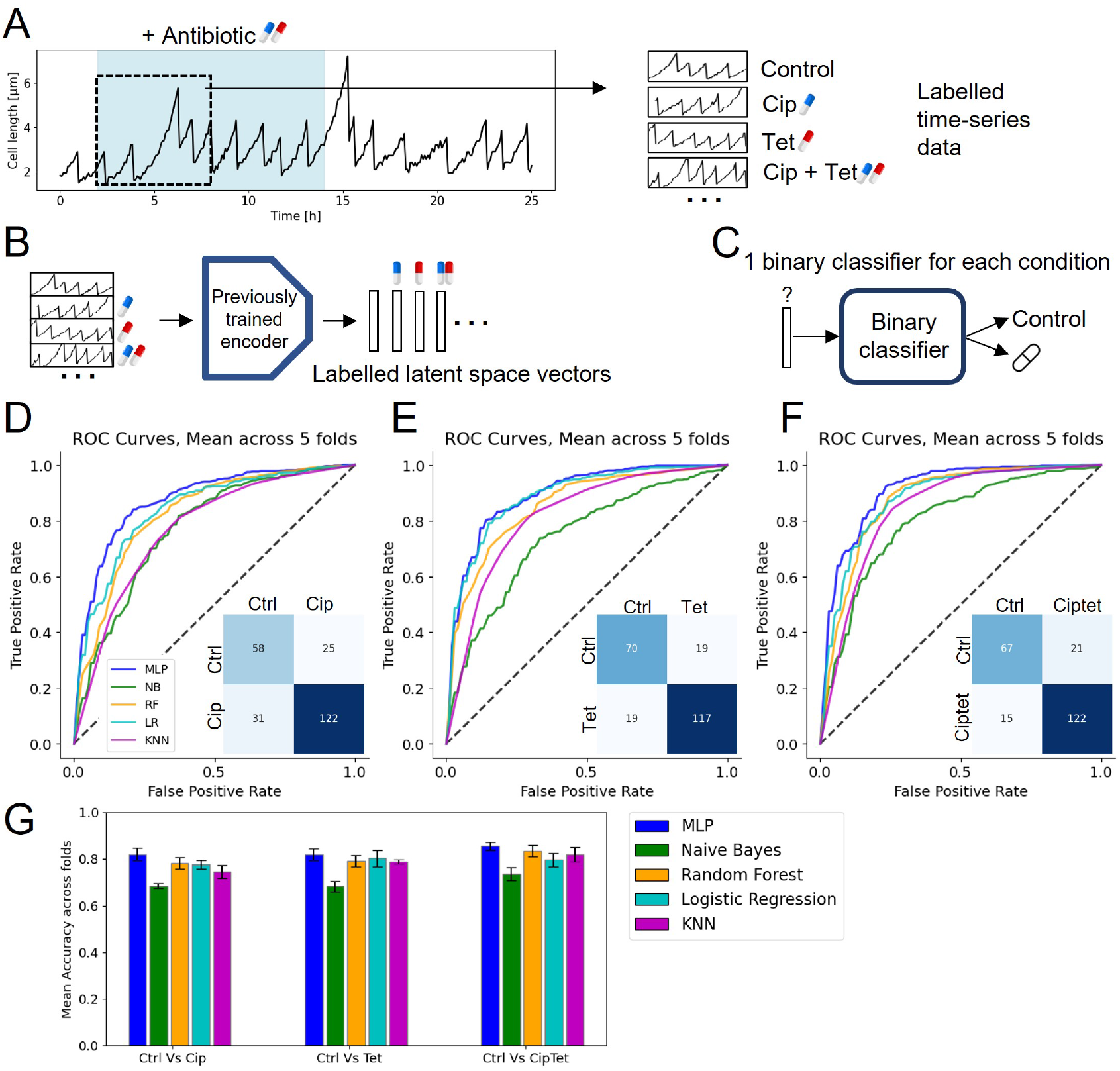
Prediction of Antibiotic Exposure Using Embedded Data. **A**: Example trajectory showing the antibiotic exposure period in blue and the sequence we used for the classification (from hour 2 to 8). We use one 6-hour time serie labelled by antibiotic condition for each mother cell. **B**: The sequences are embedded using the encoder previously trained on control data. **C**: For each antibiotic, a Binary classifier is then trained to predict whether the sequence is from a control or antibiotic exposure experiment. **D-F**: classification results on three binary tasks (Control VS Cip, Control VS Tet, and Control VS Cip+Tet), with nested 5-fold cross-validation. Mean ROC curves are shown, along with a confusion matrix created using the best-performing-fold MLP classifier and a 50% decision threshold. Results for each training fold are shown in Supplementary Figure S2. **G**: Mean classification accuracy across the 5 folds of various classifiers on the three tasks using a 50% decision threshold. All models are from sklearn: MLP = Multi Layer Perceptron (2 layers of 20 and 10 neurons), NB = Naive Bayes, RF = Random Forest classifier (50 estimators), LR = Logistic Regression, KNN = K-Nearest Neighbours classifier (3 neighbours).

We trained binary classifiers using the embeddings as input features to identify whether a time-series originates from a cell growing in normal conditions or under antibiotic exposure (Figure 2C). We focused on three binary classification tasks: control vs cip, control vs tet, and control vs ciptet. We trialled five classification models of varied complexity: Naive Bayes, Logistic Regression, a Random Forest classifier, a K-Nearest Neighbours classifier, and a Multi-Layer Perceptron. We trained the classifiers in 5-fold cross-validation on 80% of the data, totalling around 900 encoded time-series for training and 230 time-series held out as a test set for each antibiotic condition (see Table I). For ease of comparison, model hyperparameters were kept constant across the three tasks. In the three cases, labels were well balanced between the negative (control) and positive (cip, tet, or ciptet) classes. We observed promising classification performance as quantified by the Receiver Operating Characteristic (ROC) curve (Figure 2D–F). The MLP classifier displayed the best performance, achieving average AUC-ROC scores near 90% in the three tasks (Supplementary Figure S2D). Even the worst performing models were able to deliver predictions well above the naive baseline with AUC-ROC scores above 75%.

As an additional performance evaluation, we queried the best-performing MLP on the held-out test sets for the three tasks. The confusion matrices (Figure 2D–F) demonstrate good performance, with misclassification errors being balanced across false positives and false negatives, likely due to the class balance. The MLP model also displayed the best test classification accuracy (above 80%, Figure 2G) and negligible overfitting. This performance can likely be improved with further task-specific optimisation of the MLP architecture. Overall, these results strongly suggest that the VRAE embeddings have good out-of-distribution generalisation and can be paired with experimental labels for supervised prediction tasks.

### C. Prediction for external data in different growth conditions

To demonstrate the robustness of our model and its ability to transfer knowledge to prediction tasks to external datasets, we challenged it with time-series measurements from the literature. We employed cell length data from Tanouchi *et al* ^37^, consisting of *E. coli* cells growing under different temperatures. This dataset was independently generated by another laboratory using a similar microfluidic device but a different image analysis algorithm to extract the cell trajectories.

As a test of the reconstruction power of the autoencoder, we first fed the new timeseries through the encoder and decoder trained on our in-house data, which were acquired at constant temperature (37°C)(Figure 3A). The results show a remarkable reconstruction accuracy across the three growth temperatures (Figure 3B), without the need to retrain the autoencoder. The reconstructed time-series are faithful for both the normal *E. coli* growth temperature (37°C) and the lower temperatures where *E. coli* displays slower growth rates (25°C and 27°C), with a reconstruction error comparable to that observed on the test set during training. This promising result suggests a strong generalization capability of the autoencoder.

**FIG. 3.**
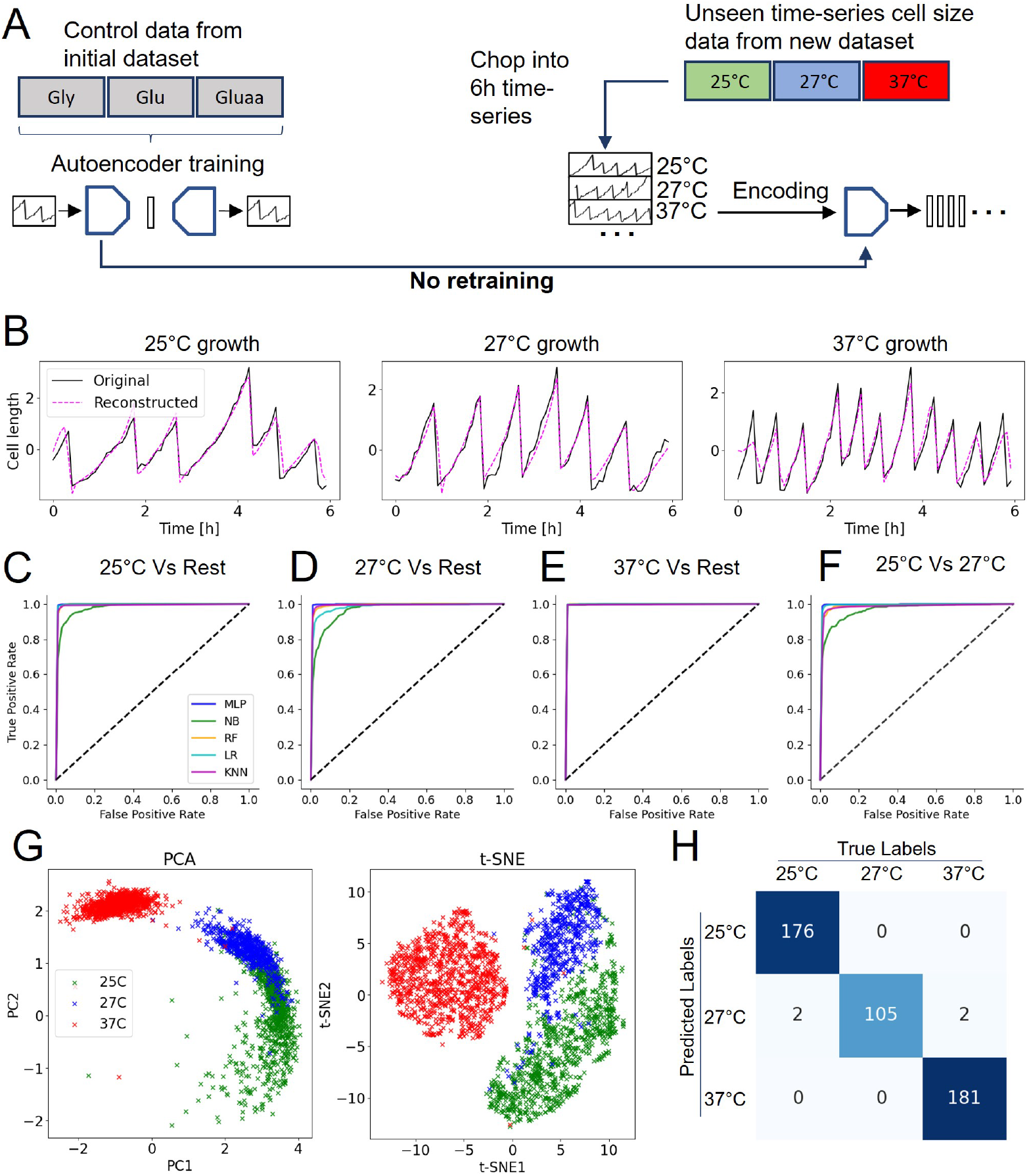
Prediction of Growth Conditions from Unseen Data Acquired by Different Laboratories. **A**: The model trained on control data on our dataset is used as it is to reconstruct and encode size time-series data from37. **B**: Sequence reconstructions by the 12-dimensional latent space autoencoder using sequences from different growth temperatures. **C–F**: Classification results of encoded data on four binary tasks. MLP = Multi Layer Perceptron (2 layers of 20 and 10 neurons), NN = Naive Bayes, RF = Random Forest classifier (50 estimators), LR = Logistic Regression, KNN = K-Nearest Neighbours classifier (3 neighbours). **G**: PCA and t-SNE representations of trajectories from the new dataset after being embedded using the autoencoder, colored by growth temperature. **H**: Confusion matrix of a 3-class classification of the three temperature groups using a Multi Layer Perceptron.

We then tested the utility of the pretrained embeddings to classify the temperature labels. We encoded the data, creating a dataset of time-series labelled by their growth temperature, and trained various binary classifiers to distinguish between different growth conditions. Figure 3C–F shows the results for different classification tasks and models. All classifiers except for Naive Bayes were able to distinguish 25°C from the other conditions with near-perfect accuracy, and with similar results for 27°C. The 37°C condition appeared even more distinct from the other in the latent space, as expected since *E. coli* grows optimally at that temperature, and a drop to 27°C or 25°C will likely have a strong effect on the elongation and division pattern. The 25°C and 27°C conditions were, as expected, harder to separate, but the classifiers were still able to distinguish them (Figure 3F) with high accuracy. The 2D projections of the latent space in Figure 3G further show the clustering of the different temperature conditions, which highlights a clear separation between 37°C and the other two conditions. We also trialled a 3-class MLP classifier with excellent predictive accuracy (Figure 3H, details in Methods). These results suggest that the autoencoder can reconstruct unseen data from other laboratories obtained in different conditions and generate meaningful representations, without the need for retraining or fine-tuning.

## III. DISCUSSION

Here, we demonstrated the use of deep representation learning for the analysis of singlecell time-series data. Using a variational recurrent autoencoder, we showed that cell trajectories from a mother machine microfluidic device can be faithfully compressed into a low-dimensional representation. These embeddings extract meaningful information with excellent generalisation performance for unseen time-series paired with experimental labels acquired in different conditions and laboratories. By projecting a large number of timeseries onto a time-independent feature space, the models can be used to qualitatively compare across experiments, for example, via well-adapted techniques such as PCA or t-SNE visualisations. Similarly, clustering algorithms can be employed to quantitatively identify relevant subgroups or outliers of biological interest. A particularly promising application is the analysis across datasets from different laboratories, for example, by pretraining models on existing data from different sources and using them for forward experimental design and generation of testable hypotheses.

Our generalisation tests focused on chemical perturbations (antibiotics) and different growth conditions (temperature). The strong generalisation performance in both cases suggests that the representations contain meaningful information about the elongation and division patterns of bacterial cells observed in different conditions. Crucially, we observed that the autoencoder can detect the antibiotic effect on cell length trajectories even when trained exclusively on data where bacteria were not exposed to the drugs. In the case of tetracycline, this strong performance somewhat agrees with expectation, as the mechanism of action of tetracycline is based on impairing protein synthesis, which is known to correlate with growth rate^38,39^. Since the autoencoder was trained on trajectories acquired in slow and fast growth, it is plausible that the training data resembles the effect of tetracycline on the temporal growth patterns. In the case of ciprofloxacin, however, the explanation is less clear because drug exposure induces DNA damage and is not directly coupled to changes in growth rate^26^. DNA damage leads to the activation of the SOS pathway and arrest of cell division while DNA is being repaired, which in turn causes cells to grow longer than their normal size. This anomaly is reflected in the temporal cell length trajectories, which is likely to be the information captured by the autoencoder.

In this work, we focused on cell length trajectories primarily because they are a baseline readout available in many mother machine experiments. The excellent predictive power of the autoencoders trained on cell length lays the groundwork for the application of this technology to other readouts that are noisier and less regular than cell length trajectories. Examples of these include fluorescence-based transcriptional reporters of stress inductio^16,28^, as well as the dynamics of appearance of fluorescent structures that report on a specific cellular event, such as the initiation of DNA replication^40^ or the emergence of mutations^18^. This type of autoencoder could also be easily scaled to handle multivariate data, for example, by combining cell length data with fluorescence reporters. A key challenge in these future applications is to endow the models with explainability power. Unlike traditional techniques for time-series analysis^4,5^, the learned embeddings are not explainable, and this can limit their use in biological discovery, particularly when we need to correlate specific latent features with a biological readout of interest. We envision that unsupervised feature extraction, together with task-specific time-series features, can be integrated into explainable and generalizable models for downstream tasks.

## IV. MATERIAL AND METHODS

### A. Single Cell time-series Datasets

#### Growth media and antibiotics dataset

We used a dataset acquired in our previous study^28^, in which *Escherichia coli* cells were trapped in the microchannels of a mother machine device^41^, pictured in Figure 1A, and perfused with growth media during the duration of the experiment. Three different growth media were tested: M9+glycerol, a minimum medium where the growth rate is low, which we refer to as “gly”, M9+glucose, an intermediate medium which we refer to as “glu”, and M9+glucose+amino acids, a richer medium where bacteria grow fast, which we refer to as “gluaa”.

Additionally, some experiments included a 12-hour exposure period to an antibiotic, from 2 hours to 14 hours of the time-series (see Figure 2A). Four different conditions were used: control (no antibiotics), ciprofloxacin (3 ng/ml), tetracycline (0.2 µg/ml), or a combination of the 2 (ciprofloxacin three ng/ml + tetracycline 0.2 µg/ml), which we refer to as “ciptet”. Table S1 shows the list of experiments and how many single-cell time-series are available in each one.

Images were taken every 5 minutes (10 in gly) to generate timelapses of the cells in their microchannels. The images were processed using the BACMANN software^20^ to segment individual cells, track them and extract the time-series. The time-series data used in this study were obtained by monitoring only the mother cells, which is are cells located at the very top of the channel (see Figure 1A). As every daughter cell gets ultimately pushed out of the channel by the mother cell’s divisions, the mother cell is the only cell which can be followed for extended periods of time.

For our work, we used the trajectories corresponding to the evolution of the mother cell’s length (a proxy for cell volume as the width is constant) over time. Since bacteria constantly grow and divide, these trajectories follow a saw-tooth pattern of elongation followed by a sudden drop as the cell splits in half to create a daughter cell, as shown in the example in Figure 1B. Each trajectory is 275 to 300 time points long (with a portion of the trajectories ending during the experiment as the analysis software loses track of certain cells). Glycerol time-series were adjusted to have time points every 5 minutes by filling the gaps using the mean of adjacent points. time-series that ended before the 14-hour mark were removed, as well as time-series for which the cell length exceeded the channel length of 20 µm, since part of the cell would effectively be off-camera.

#### Temperatures dataset

The dataset was retrieved from the study by Tanouchi *et al* ^37^. The experimental setup is similar to the one presented in the dataset above. A mother machine and *E. coli* bacteria were also used. The media used for all experiments was LB. In this dataset, the cells were grown at three different temperatures: 25°C, 27°C, and their preferred growth temperature, 37°C. The dataset contains 65 25°C time-series, 54 27°C time-series, and 160 37°C time-series. Images were taken every minute; In our analysis, the time-series were downscaled to one time point every five minutes to match the time-series from the first dataset.

### B. Training a Variational Recurrent autoencoder for time-series

#### Training set and data augmentation

We composed a training dataset combining one replica of each of the three control experiments from the gly, glu and gluaa media (see Table S1). The time-series were randomly shuffled and 10% was set aside as a test set. We decided to train the model on control (no antibiotic) data from all three growth media, using the first 14 hours of each trajectory to limit age-related cell death^10,42^. In the rare event that a cell dies during the experiment, the time-series was cropped immediately before cell death. We used the time of death provided with the dataset. The time-series were then normalised (zero-mean, unit variance, normal distribution) using the tslearn Python module.

In the training set, the time-series were chopped into 72-time-point (6-hour) sequences. We used a rolling window of 6 time points (30 minutes), which created a set of overlapping 6-hour fragments, 6 time points (30 minutes apart) apart. This artificially increased the content of our training set as a form of data augmentation. The training set had a total of 57607 6-hour time-series (see Table I). In the test set, we also used 72 time-point sequences, but they did not overlap, for a final number of 699 time-series (see Table I).

#### Autoencoder architecture and training

The autoencoder is a variational recurrent autoencoder with both the encoder and decoder being LSTM models. Its architecture was retrieved from a previous study (^23^), which successfully trained it on temporal data. For a regular autoencoder, this Bayesian approach maps the data to a latent space; however, in this case, the data is mapped to a distribution over latent variables, as illustrated in Figure S1 and^23^.

We used two hidden layers, each with 90 units, for both the encoder and the decoder. The training was done in Python using PyTorch. We built 19 model architectures with varying latent space sizes: from 2 to 20. Each of those architectures was trained 10 times, for 200 epochs. We used the Adam optimiser with a learning rate of 0.001, and used a dropout rate of 0.2 to reduce overfitting. The loss function used was a hybrid loss that combined a reconstruction loss, which compares the input and output, with a Kullback-Leibler (KL) divergence loss. This KL loss encourages the learned latent variable distribution to be close to a standard normal distribution and is necessary for training variational autoencoders^23^. After every epoch, the models were used to reconstruct all the trajectories from the test set and the total reconstruction loss was recorded. This test set reconstruction loss for the different latent lengths is shown in Figure 1D, along with reconstruction examples.

### C. Choosing the Latent Space size hyperparameter Using Clustering

To determine the optimal latent space size, we evaluated how well the three growth conditions are clustered in the latent space, as a measure of representation quality. We performed clustering on the total dataset of control experiments (i.e. the data used for training, as well as additional experiments not seen by the trained model). All the time-series were normalised and chopped into non-overlapping 6-hour sequences, then transformed using the encoder half of one of the models. The k-means algorithm (sklearn Python package) was then used to cluster this embedded data into 3 clusters (using 100 random initialisations).

To evaluate the clustering performance of each of the 19 autoencoders, we calculated the Adjusted Rand Index^43^ between the predicted clusters and the three known growth conditions labels (gly, glu and glu+aa). It ranges between 0 and 1 and assesses how well the three clusters correspond to the three growth conditions, adjusting for chance. This measurement was used in conjunction with the test set final reconstruction loss to choose the Latent Space Size of 12 in Figure 1E.

### D. Classification using embedded time-series

Classification was performed using a variety of classifier models: Naive Bayes, Logistic Regression, a Random Forest classifier with 50 estimators, a K-Nearest Neighbours (KNN) classifier (3 neighbours) and a simple Multi Layer Perceptron (2 layers of 20 and 10 neurons). Models were fit using their Python SKlearn implementation. Figure 2 models are binary classifiers trained to take as input a latent vector from an encoded time-series and predict either 0 (the time-series comes from a control experiment) or 1 (the time-series comes from an Antibiotic experiment) as presented in Figure 2B-C. In Figure 3H, the 3-class classifier used was a Multi Layer Perceptron with two layers of 20 and 10 neurons (sklearn).

## ACKNOWLEDGEMENTS

A. F. was supported by the United Kingdom Research and Innovation (EP/S02431X/1), UKRI Centre for Doctoral Training in Biomedical AI at the University of Edinburgh.

M. E. K. was supported by a Wellcome Trust Investigator Award (Grant No. 205008/Z/16/Z). We thank James Broughton for assistance with data generation and analysis. Portions of the manuscript text were edited using Grammarly, a language assistance tool. The authors reviewed and approved all AI-assisted content.

## DATA AND CODE AVAILABILITY

The datasets employed in this study are publicly available in Zenodo^44^ and Figshare^29^ repositories. Code for training the VRAE architecture can be found in https://github.com/tejaslodaya/timeseries-clustering-vae. Our code for model training and generating the paper figures has been deposited in Zenodo^45^.

## SUPPLEMENTARY INFORMATION

**Supplementary Figure S1.**
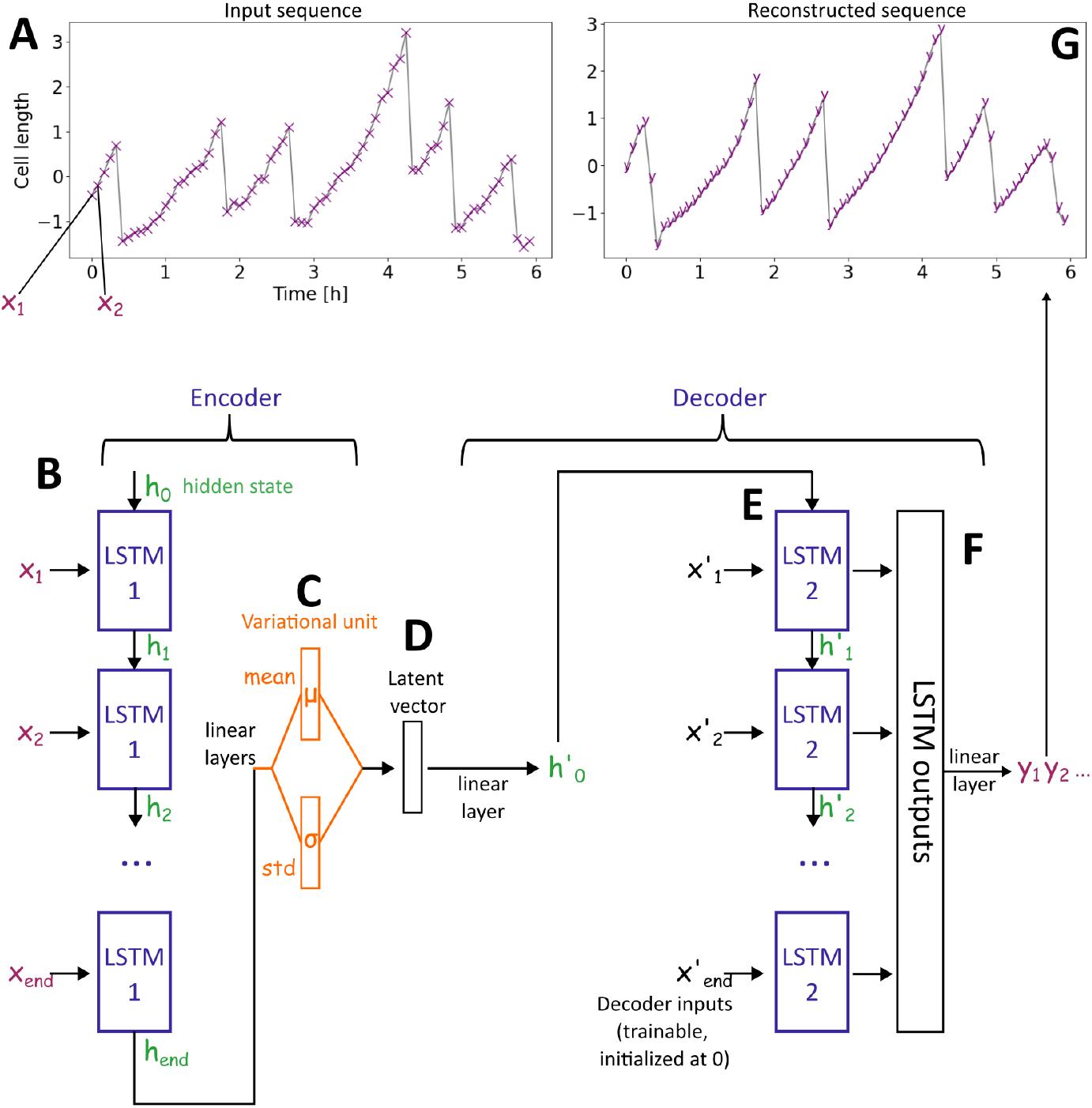
Architecture of the Variational Recurrent autoencoder. The 72 timepoints *X*_*i*_ composing the input sequence **A** are passed through the first LSTM model **B** one by one in temporal order. The hidden state of the LSTM *h* is updated with each iteration. After passing all the timepoints through, the final hidden state *h*_*end*_ is transformed by two different linear layers into two vectors of size equal to the size of the latent vector. **C**: A vector of means and a vector of standard deviations, one for each latent feature. During training, the latent vector **D** is generated by sampling from these normal distributions; outside of training, the vector of means is used as the latent vector. The whole part until this point is called the Encoder, whose output is the latent vector. The Decoder begins with a linear layer generating a hidden state vector *h*^*′*^_0_ to initialise the second LSTM **E**. The second LSTM inputs are initialised at 0 and trained along with the rest of the model. A linear layer finally transforms the outputs of this second LSTM to compose the output sequence *y*_*i*_.

**Supplementary Figure S2.**
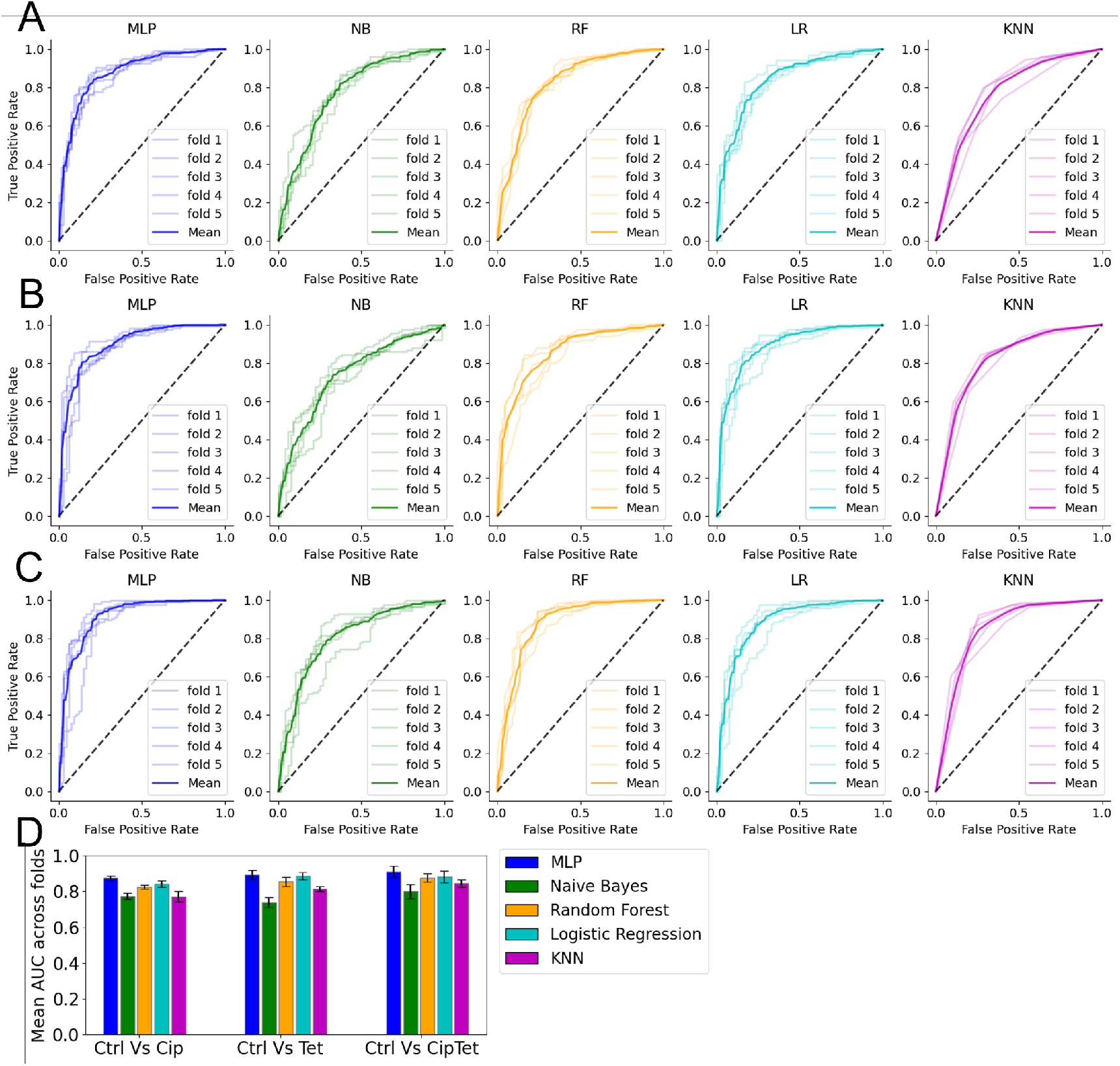
Cross validation ROC curves and AUCs for control vs antibiotic classification. **A-C**: ROC curves for all 5 folds (Mean in bold) using different classifier architectures. **A**: Control vs Cip, **B**: Control vs Tet, **C**: Control vs Ciptet. MLP = Multi Layer Perceptron (2 layers of 20 and 10 neurons), NB = Naive Bayes, RF = Random Forest classifier (50 estimators), LR = Logistic Regression, KNN = K-Nearest Neighbours classifier (3 neighbours). **D**: Mean Area Under the Curve for each task and classifier over the 5 folds. Error bars show standard deviation.

**Supplementary Table S1.** Number of mother cell time-series available in each experiment and replicate from the antibiotics dataset. Control = no antibiotic, Cip = ciprofloxacin (3 ng/mL), Tet = tetracycline (0.2 µg/mL), or a Cip+Tet = ciprofloxacin 3 ng/ml + tetracycline 0.2 µg/mL, Gly = M9+glycerol, Glu = M9+glucose, Gluaa = M9+glucose+amino acids.

|  | Control | Cip | Tet | Cip+Tet |
| --- | --- | --- | --- | --- |
| Gly | Replicate 1: 188 | Replicate 1: 252<br>Replicate 2: 307 | Replicate 1: 317 | Replicate 1: 328 |
|  | Replicate 2: 376 |  | Replicate 2: 277 | Replicate 2: 431 |
|  | Replicate 3 :304 |  | Replicate 3 :411 |  |
| Glu | Replicate 1: 78 | Replicate 1: 386 | Replicate 1: 351 | Replicate 1: 355 |
|  | Replicate 2: 342 | Replicate 2: 377 | Replicate 2: 354 | Replicate 2: 350 |
| Gluaa | Replicate 1: 283 | Replicate 1: 218 | Replicate 1: 327<br>Replicate 2: 237 | Replicate 1: 343 |
|  | Replicate 2: 334 | Replicate 2: 191 |  | Replicate 2: 304 |
|  |  | Replicate 3 :235 |  |  |

